# Decoding human placental cellular and molecular responses to obesity and fetal growth

**DOI:** 10.1101/2025.01.27.634981

**Authors:** Hong Jiang, Emilie Derisoud, Denise Parreira, Nayere Taebnia, Paulo R. Jannig, Reza Zandi Shafagh, Allan Zhao, Congru Li, Macarena Ortiz, Manuel Alejandro Maliqueo, Elisabet Stener-Victorin, Volker M. Lauschke, Qiaolin Deng

**Author notes:** These authors have contributed equally to this work and share first authorship (Hong Jiang, Emilie Derisoud).

## Abstract

Maternal obesity increases risk of large-for-gestational-age (LGA) births and subsequent cardiometabolic disorders in offspring. To identify placental signatures associated with these outcomes, we performed single-nucleus RNAseq on placentas from women with obesity delivering appropriate-for-gestational-age (AGA) or LGA infants, compared to normal-weight controls. In maternal obesity, regardless of fetal growth, syncytiotrophoblasts showed upregulated hypoxia and TNF-α signaling, while cytotrophoblasts exhibited downregulated receptor tyrosine kinase signaling. However, only in LGA placentas, villous non-trophoblasts displayed upregulated TNF-α signaling and inflammatory responses. Notably, Hofbauer cells in LGA placentas presented transcriptional alterations in immunometabolism-related genes and functioned as main signaling senders via *SPP1*. Importantly, we recapitulated syncytiotrophoblast responses to maternal obesity using a novel microfluidic organoids-on-a-chip co-culture. These findings reveal distinct transcriptional responses of placental cell types to maternal obesity and fetal growth, highlighting potential intervention pathways to mitigate future disease risks.

## Introduction

The global prevalence of obesity among women of reproductive age reached 18.5% in 2022, making maternal obesity an increasing concern for women’s health (*1*). Maternal obesity is manifested by chronic low-grade inflammation, hyperinsulinemia, and dysregulated lipid metabolism (*2*). Consequently, these systemic changes collectively impact the intrauterine environment and lead to immediate and long-term health consequences for both mothers and their offspring including greater risk to develop cardiometabolic diseases and type 2 diabetes (*3–5*). Among pregnancy complications, delivery of large-for-gestational-age (LGA) infants (birthweight > 90^th^ percentile (*6*)) is one of the most frequently observed ones, occurring in 13.41%-38.3% of pregnancies affected by obesity (*7–10*). Notably, LGA itself is not only a pregnancy complication but also a critical risk factor for further adverse outcomes, including birth trauma (*11*), childhood obesity (*12*), cardiometabolic and neurological disorders in early adulthood (*13*). Therefore, preventing LGA may serve as an effective strategy to mitigate the cascade of associated complications.

Maternal obesity is often associated with excessive gestational weight gain (*14*), which subsequently contributes to a higher risk of LGA infants due to excess nutrient supply (*15*). However, the UK UPBEAT study, a randomized controlled trial testing a tailored complex lifestyle intervention (diet and physical activity) in pregnant women with obesity, found that reduction in gestational weight gain did not have significant effects on the outcomes of gestational diabetes and LGA infants (*16*). Also, the EMPOWaR study found that pharmacological use of metformin in pregnant women with obesity had no significant effect on reducing birthweight percentiles or preventing adverse neonatal outcomes despite clearly improving maternal metabolic health (*17*). These studies collectively pinpoint the potential additional mechanisms contributing to fetal growth, and lack of success in preventing LGA infants with current interventions hence underscores the need for a deeper understanding of the underlying mechanisms.

Intriguingly, not all fetuses exposed to maternal obesity exhibit excessive growth and develop LGA, with some still following a normative growth trajectory resulting in AGA. This intrinsic variability may reflect an inherent adaptive mechanism *in utero* that could be harnessed for future LGA management. The placenta is the central structure at the maternal-fetal interface, regulating the quantity and quality of nutrient supply from the maternal circulation by adjusting its transport capacity and nutrient uptake (*18*). Additionally, it dynamically modulates growth factors and hormonal secretion (*19*) and adjusts inflammation levels (*20*) to align with the growth requirements of the fetus. These activities are intertwined and could serve as the link between maternal obesity and divergent fetal growth trajectories. For example, elevated maternal insulin levels in obesity activate mTOR signaling as well as glucose and amino acid transport in the placenta, thus contributing to accelerated fetal growth in some women with obesity (*21*). Circulating pro-inflammatory cytokines in obesity, such as IL-6, tumor necrosis factor-α (TNF-α) and leptin are found to promote lipid storage in the placenta (*20*) and enhance amino acid and lipid transporters in trophoblast. Moreover, secretion of placental lactogen and prolactin are correlated with pre-pregnancy BMI and fetal growth (*22*), further supporting the active role of the placenta in modulating the effects of obesity on fetal growth (*23*).

However, current understanding of the placental response to maternal obesity largely relies on gene expression analyses conducted at the whole-tissue level or in cell cultures (*24–27*), which are often confounded by tissue complexity or lack of physiological relevance in culture conditions. Therefore, identifying molecular signatures in the placenta associated with AGA and LGA infants at a single-cell level may provide valuable insights into cell-type specific mechanisms driving distinct growth patterns and inform potential targeted therapeutic strategies. Here, we performed single-nucleus RNAseq (snRNA-seq) on term placentas collected in a Chilean cohort study, in which women with obesity that did not develop other pregnancy complications except for more frequent delivery of LGA infants (*28*). We divided the placental samples and analyzed three groups: normal BMI with AGA infants (Control), maternal obesity with AGA infants (O-A), and maternal obesity with LGA infants (O-L). Cell-type specific transcriptomic profiles and pathways were identified to be either share or different in O-A and O-L groups, suggesting molecular responses to maternal obesity or fetal growth, respectively. Moreover, the intercellular ligand-receptor communication network revealed the major cell types mediating these maternal or fetal effects. Finally, microfluidic co-culture of adipose spheroids and trophoblast organoids could recapitulate some transcriptional changes of trophoblasts in maternal obesity, providing an *in vitro* model with mechanistic insights to study the effects of maternal conditions on human placentas in future.

## Results

### Cell types in placentas stratified by maternal obesity and fetal growth

To analyze the placental transcriptional changes in maternal obesity and investigate the interplay between the placenta and fetal overgrowth, we selected 12 samples from a birth cohort with representative maternal BMI and birthweights for snRNA-seq (*28*) (Fig. 1A and fig. S1A). All samples were from vaginal delivery and the O-A and O-L groups had no significant differences in maternal age, placental weight and placental efficiency compared with Controls (Fig. 1B, Fig. 1A). We dissected the placental villous region and dissociated the tissue into single nuclei, followed by library preparation using 10x Chromium (Fig. 1C). A total of 37,408 nuclei were retained for downstream analyses after stringent quality control, yielding comparable total counts and gene number per cell across all groups (Fig. 1D). Using canonical marker genes for cell types (*29*), we annotated 14 placental cell types, including five types of trophoblasts, two states of endothelial cells, two types of fibroblasts, and five types of leukocytes (Fig. 1E). There was no significant difference in the proportion of cell types, either in various trophoblasts or in different villous non-trophoblast cells, across the three groups (Fig. 1F, G and Appendix Table S1).

**Fig. 1:**
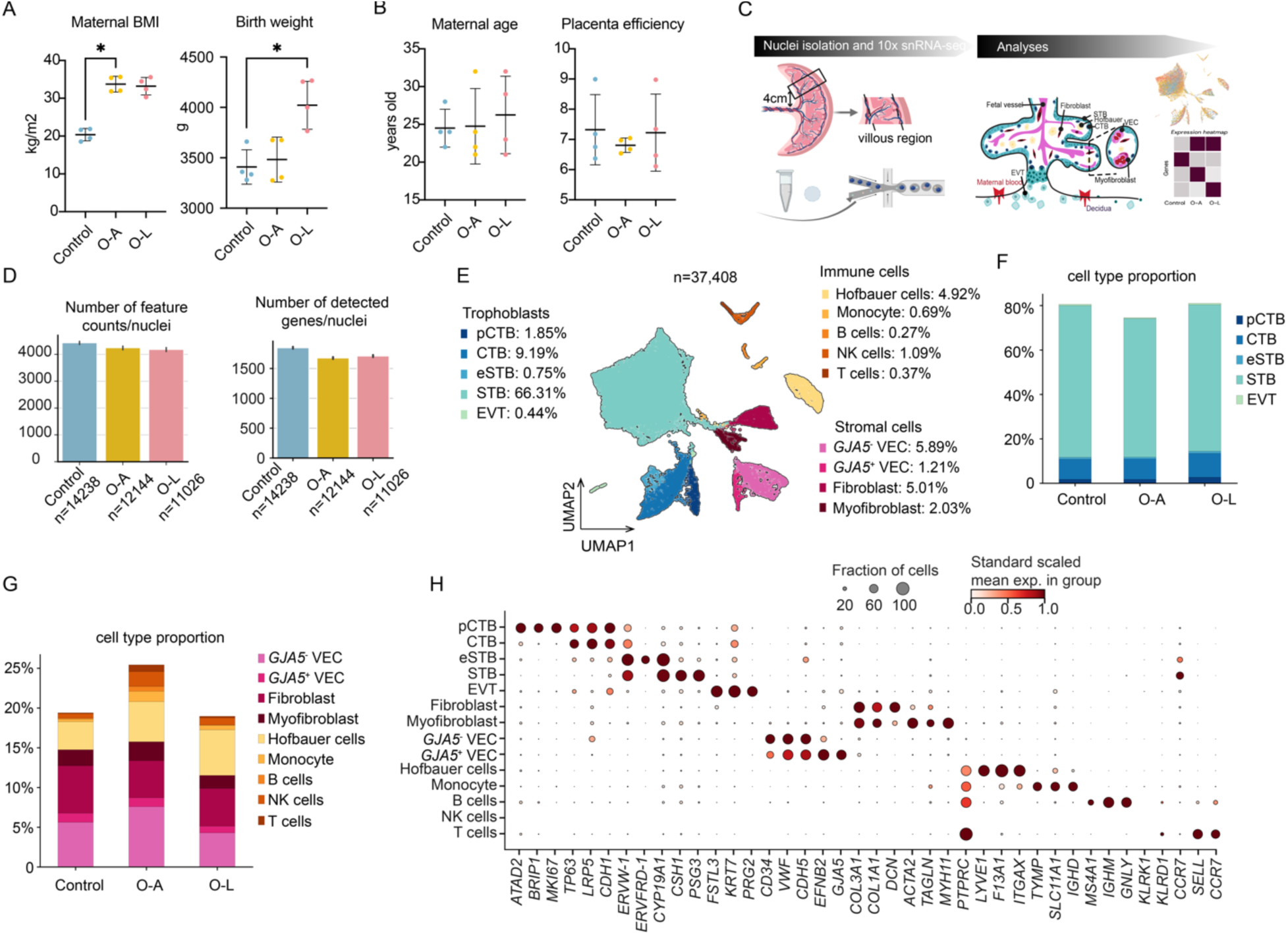
snRNA-seq of human placentas from normal-weight mothers and mothers with obesity stratified by fetal birthweight. (**A)** Maternal BMI (body mass index), and birthweight of samples for snRNA-seq from each group. **(B)** Maternal age, and placental efficiency of samples for snRNA-seq from each group. **(C)** Experimental procedures. **(D)** Number of feature counts per nuclei and number of detected genes per nuclei. **(E)** Two-dimensional UMAP (Uniform manifold approximation and projection) of 37,408 nuclei profiles (dots) from all donors (n=12), colored by cell type. (**F)** Stacked barplot showing trophoblast cell proportion in each group. **(G)** Stacked barplot showing the proportion of non-trophoblast cell types i n each group. **(H)** Mean expression (dot color, gene standard scaled across all cells of a type) and fraction of expressing nuclei (dot size) of marker genes for each of the 14 major cell types. Controls: normal-weight women (n=4); O-A: Mothers with obesity with appropriate-for-gestational-age baby (birthweight within the 25^th^ and 90^th^ percentile of WHO growth curves); O-L: Mothers with obesity with large-for-gestational-age baby (birthweight >90^th^ percentile of WHO growth curves); CTB: cytotrophoblasts; VEC: vascular endothelial cells; EVT: extravillous trophoblasts; pCTB: proliferative CTB; STB: syncytiotrophoblasts; eSTB: early syncytiotrophoblasts.

Trophoblasts comprised the majority of cells, primarily syncytiotrophoblast (STB) and villous cytotrophoblasts (CTBs). Proliferative CTBs (pCTBs) were identified as being in the cell cycle (Fig. 1H, fig. S1B). Co-expression of *ERVW-1* (Syncytin-1) and *ERVFRD-1* (Syncytin-2) identified early STB (eSTB), consistent with previous findings (*30*). Mature STB shared expression of *ERVW-* 1 and *CYP19A1* with eSTB but uniquely expressed *CSH1* and *PSG3* (Fig. 1H). The extravillous trophoblasts (EVTs) accounted for only 0.5% of all nuclei, with fewer than one hundred per group, aligning with our villous sampling approach (Appendix Table S1). Among other villous non-trophoblast cells, stromal cells within the chorionic villi included fibroblasts and myofibroblasts. Myofibroblasts were annotated by concurrently expressing collagen (*COL3A1* and *COL1A1*), actin (*ACTA2*) and myosin subunits (*MYH11*) (Fig. 1H). Vascular endothelial cells (VECs) were divided into a small subset of *GJA5*^+^ and *GJA5^−^* VEC (Fig. 1H and fig. S1C). *GJA5* is a gap junction protein facilitating direct intercellular communication by forming channels. Among the immune cells, we primarily captured Hofbauer cells, specialized macrophages located within the chorionic villi (Fig. 1H). *PTPRC* was also detected in other remaining nuclei, indicating their leukocyte origin. These remaining nuclei could be further annotated into monocyte, B cells, NK cells and T cells (Fig. 1H and Appendix Table S1). However, due to their limited numbers (fig. S1D), likely carried over from dissection, and being outside the scope of our study, these cells were excluded from subsequent analyses.

Next, we focused on the major villous cell types, i.e., STB, CTB, VEC, fibroblasts and Hofbauer cells, to understand their distinct molecular signatures in the placental responses to obesity and regulation of fetal growth. To identify functional pathways and gene expression changes in each cell type, we developed a scoring (C-score) metric to quantitatively measure the transcriptional changes in the two obesity groups, O-A and O-L compared to the control. The C-score metric was calculated based on the effect size and the false discovery rate from differential gene expression analysis and gene set enrichment analysis followed by ranking. A high positive C-score indicated maternal effects independent of fetal overgrowth, as these genes and pathways were similarly regulated in both comparisons (i.e., O-A vs. control and O-L vs. control). In contrast, a low negative C-score indicated effects linked to fetal growth effects, as the O-A and O-L groups exhibited different changes (see Methods: Scoring the shared or different DEGs between comparisons). This method provided a systematic approach to rank changes due to maternal or fetal growth effects across all cell types and cell-cell communications in the placenta, without relying on arbitrary fold-change cutoffs. The number of the shared or different DEGs was provided in Appendix Table S2.

### Identification of STB subpopulations along the maturation states

The STB layer is formed through the fusion of CTBs, a process that requires remodeling of cytoskeletal and plasma membrane dynamics. A total of 24,478 STB nuclei were captured, enabling us to investigate the subpopulations across the different phases of STB life cycle. We first re-clustered the STB nuclei and identified three substates (STB-a, -b and -c) (Fig. 2A). To better elucidate their maturation process, we incorporated CTBs and eSTBs as early differentiation timepoints to place STBs in context for pseudo-time analysis (fig. S2A). The three STB substates were aligned along the pseudo-time axis with varying numbers of cells distributed across the timeline (Fig. 2B). Novel gene markers were subsequently identified, highlighting the biological differences among the substates (Fig. 2C and fig. S2B). STB-a appeared at an earlier stage characterized by the expression of *TENM3* (Fig. 2C, fig. S2C). *TENM3* encodes a teneurin transmembrane protein that promotes homophilic cellular adhesion (*31*). *TENM3*^+^ STB was demonstrated in the STB layer by *in situ* hybridization (*32*). This state was also characterized by genes involved in cytoskeleton, tight junction, and extracellular matrix (*ITGB1*, collagen subunits *COL4A1, COL4A2*, *FN1* and *TIMP3*) (Fig. 2C). Furthermore, STB-b specifically expressed *PDE4D* (phosphodiesterase 4D), a cAMP-specific phosphodiesterases reported to limit oversyncytialization (*33*) (Fig. 2C). Finally, the mature STB-c state was characterized by the expression of tight junction inhibitors such as and *PKIB* (Fig. 2C, fig. S2C). *ADAMTS6* could inhibit cell-cell junctions and was previously identified as a marker of a distinct STB state (*34*, *35*). *PKIB* is involved in the cAMP pathway, crucial for STB formation (*33*). Accordingly, STB-b nuclei appeared to balance syncytialization, while STB-c nuclei actively engage in the process, collectively contributing to a dynamic balance. Excessive syncytialization leads to release of aged cytosolic material into maternal circulation, while restricted fusion results in exhaustion of the syncytial layer (*36*). Therefore, the dynamics of these three subpopulations may play a critical role in maintaining STB homeostasis. Reassuringly, these STB states were further confirmed and in line with previously identified various STB maturing substates by integrating our data with a recently published term placenta snRNA-seq dataset (*37*) using scANVI (*38*) (fig. S2D). The percentage of three STB states was not significantly different among three groups (Fig. 2D).

**Fig. 2:**
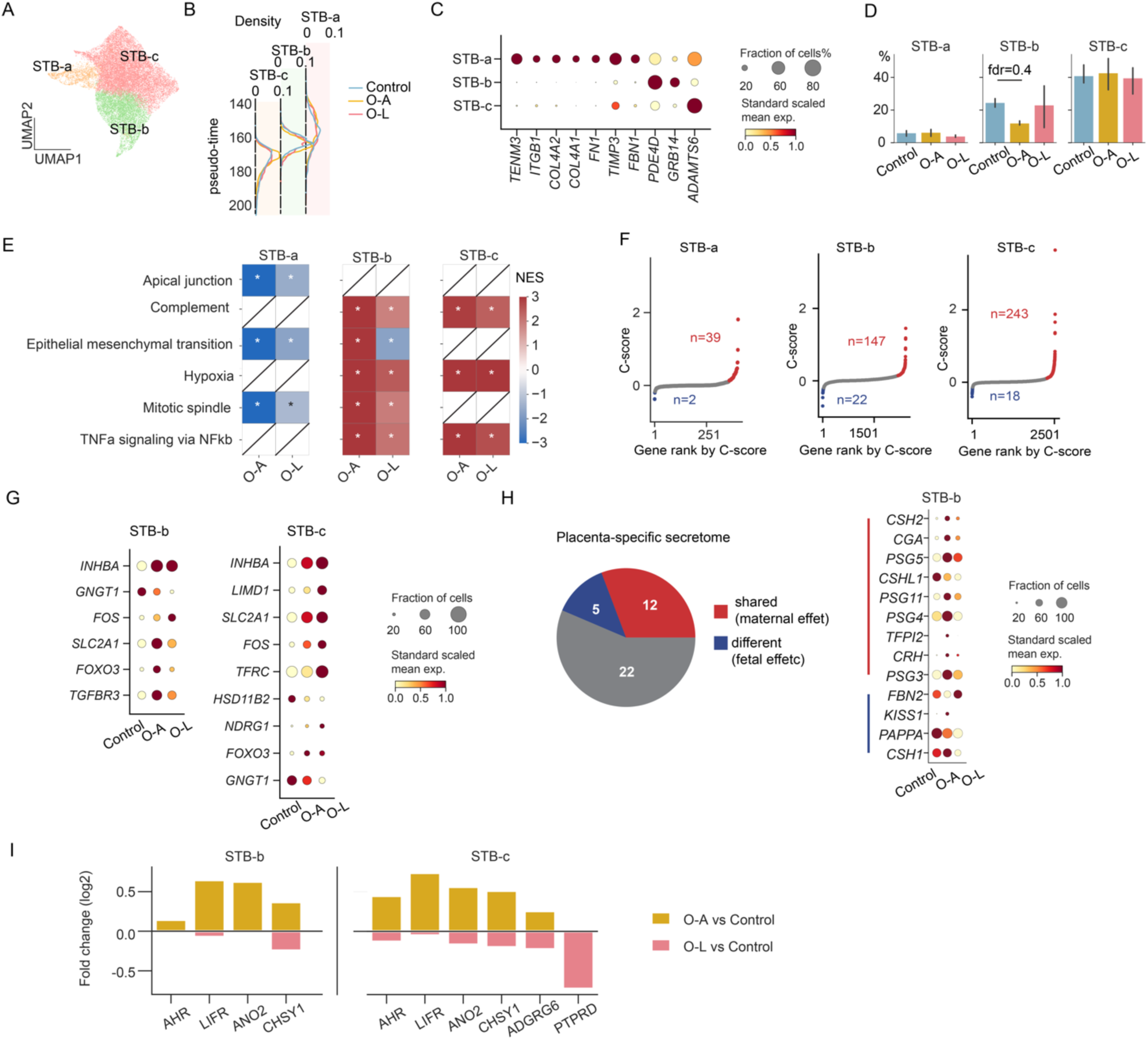
Maternal obesity changed the expressionof hypoxia-, inflammation-nutrient transport-related pathway in STB nuclei. **(A)** UMAP representation of STB colored by STB states. **(B)** Pseudotime value visualization across trophoblast cell types and states. Density plot of pseudotime value colored by each group. **(C)** Dotplot of genes specifically expressed in STB-a, -b or -c state. Dot color indicates the standard-scaled mean expression level of the gene. **(D)** Proportion of STB states across groups. Bar height represents the mean and the error bar represents 50% interval. Data were compared by a Bayesian model. **(E)** Gene set enrichment analysis (GSEA) with the enrichment score of hallmark gene sets for O-A *vs* Control and O-L *vs* Control. NES: normalized enrichment score. *adjusted p-value < 0.05. **(F)** Scatterplots of genes ordered by C-scores in STB. **(G)** Dotplots showing the expression of the shared DEGs in O-A and O-L involved in hypoxia pathway. **(H)** The proportion of shared or different DEGs in O-A and O-L among the placenta-specific secretome reported in human protein atlas (*46*) (left). Dotplot showing the expression of the genes in STB-b that encode placenta-specific secreted proteins. **(I)** Barplots showing the fold changes of DEGs related to fetal growth effects (different in O-A and O-L) encoding for receptors, channels or molecular sensors.

### Transcriptional changes due to maternal or fetal growth effects among STB nuclei

The STB nuclei at different transcriptional states possibly varies in responses to surrounding signals. We first identify the underlying biological pathways in maternal obesity with or without fetal overgrowth utilized hallmark gene sets from MSigDB (*39*). Gene set enrichment analysis (GSEA) showed that O-A and O-L shared dysregulated pathways related to epithelial mesenchymal transition (EMT), apical junction and mitotic spindle regions in STB-a nuclei (Fig. 2E and Appendix Table S3). STB-b and STB-c also shared several dysregulated pathways in both groups, including complement, hypoxia, and TNF-α signaling via NF-kβ, all of which are indicative of inflammatory processes (Fig. 2E and Appendix Table S3). To determine genes within these pathways that may contribute to the pathways, we intersected shared and different DEGs with pathway-specific gene sets focusing on the hypoxia and TNF-α signaling in STB-b and STB-c (Fig. 2F). Among those shared genes driving the enrichment of the hypoxia signaling pathway, *FOXO3* and *FOS* have been reported as the crosstalk between hypoxia and TNF-α signaling (*40–43*). Important transporters such as *SLC2A1* (i.e. GLUT1) and *TFRC* may mediate the interaction between hypoxia, nutrient transfer and metabolism (*44, 45*) (Fig. 2G). *SLC2A1* is predominately expressed in the STB to facilitate glucose uptake from the maternal circulation and *TFRC* mediates cellular iron uptake. Expression of both is shown to be regulated by hypoxia (*45*).

Next, we used a previously published set of placenta-specific secreted proteins from the human secretome project (*46*). A total of 12 genes were identified being altered by maternal obesity, among which most were upregulated except for *CSHL1* (Fig. 2H). *CGA* encodes the alpha unit of human glycoprotein hormones, including hCG (chorionic gonadotropin). Changes in hCG levels have been linked to adverse pregnancy outcomes, and its dysregulation highlights the detrimental effects of maternal obesity on the placenta (*47*). A total of 5 genes encoding placenta-specific secreted proteins were identified to be regulated according to fetal growth. Specifically, *FBN2* was upregulated while *KISS1*, *PAPPA* and *CSH1* were downregulated in O-L (Fig. 2H). Notably, *FBN2* encodes placensin, a glucogenic hormone highly expressed in human placentas to stimulate hepatic glucose secretion and cAMP production (*48*). It was elevated only in O-L but decreased in O-A, in line with its increased expression in pregnancy complications such as gestational diabetes. *PAPPA* encodes a metalloprotease secreted by the human placenta that modulates IGF bioavailability. Its concentrations are inversely correlated with glycemia and odds of developing gestational diabetes (*49*), aligning with our finding of the greatest reduction observed in the O-L group (Fig 2H). Such different hormone regulation resulted from the placental responses to maternal uterine conditions likely contributed to the different fetal growth effects.

Furthermore, the expression of receptors or channels that sense the maternal metabolic signals could also be altered accordingly to fetal growth. We found that a total of 7 genes being altered differently between O-A and O-L (Fig. 2I). Among those expressed in both STB-b and STB-c, *AHR* encodes the aryl hydrocarbon receptor that senses the varied cellular environment, including endogenous tryptophan derivatives, an essential amino acid critical for placental and fetal development (*50*). *LIFR* encodes the receptor of leukemia inhibitory factor, and LIFR deficiency leads to impaired placenta differentiation, reduced embryo viability, and increased birth weight in mice (*51*). *ANO2* encodes a calcium-gated chloride channel and *CHSY1* is an enzyme for synthesis of chondroitin sulfate in response to nutrient levels. All these genes were only upregulated in O-A but not in O-L, suggesting an inadequate response in O-L (Fig. 2I).

### PI3K-AKT and MAPK cascade signaling pathways changes in CTBs

Next, we analyzed the enrichment of hallmark gene sets in CTBs, the cell type that positions beneath STB. In contrast to STB, the hallmark gene sets were we predominantly decreased in both O-A and O-L (Appendix Table S3). These included apical junction, mitotic spindle, G2M checkpoint, heme metabolism, hypoxia, IL2-STAT5 signaling (Fig. 3A). In contrast, Myc targets showed a divergent response with downregulation only in O-L (Fig. 3A). MYC target pathway, critical for CTB proliferation, differentiation, and metabolic adaptation, is often suppressed by inflammation and hypoxia (*52*).

**Fig. 3:**
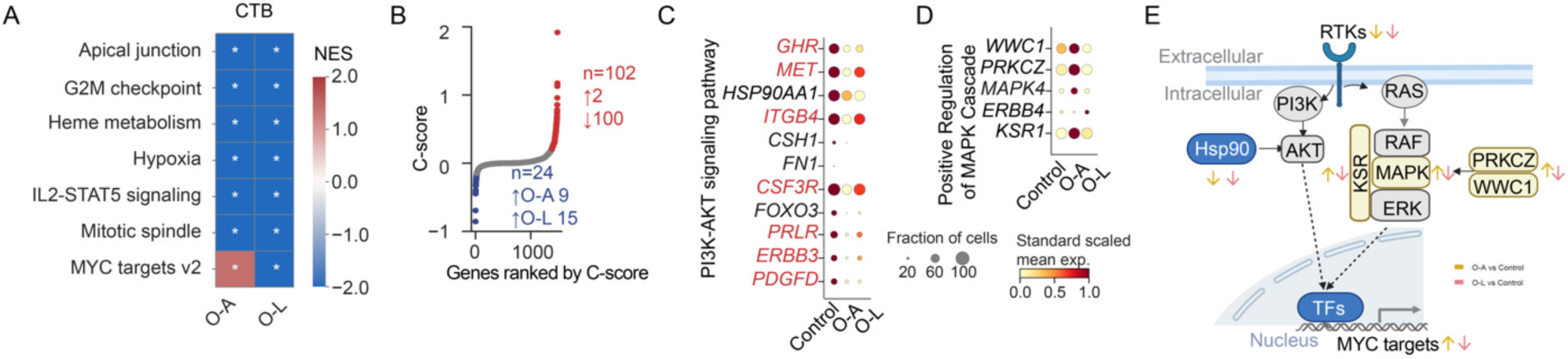
Gene expression changesin PI3K-Akt pathway and MAPK cascade in CTBs from mothers with obesity with or without fetal overgrowth. **(A)** Gene set enrichment analysis (GSEA) with the enrichment score of hallmark gene sets for O-A *vs* Control and O-L *vs* Control. NES: normalized enrichment score. *indicates adjusted p-value < 0.05. **(B)** Scatterplots of genes ordered by C-scores in CTB. **(C)** Dotplots showing the expression of the genes involved in PI3K-Akt signaling pathway, and **(D)** positive regulation of MAPK cascade. **(E)** Summary of genes involved in the pathways downstream the receptor tyrosin kinase (RTK) signaling. The PI3K-Akt signaling was reduced in CTBs from both obesity groups, while the MAPK signaling was upregulated in O-A.

Next, we identified 102 genes as altered by maternal obesity in both O-A and O-L, among which 100 genes were downregulated genes (Fig 3B). Interestingly, the altered genes were significantly enriched in the PI3K-Akt signaling pathway, with those highlighted in red specifically involved in receptor tyrosine kinase (RTK)-mediated transduction of growth factor signals (Fig. 3C). In addition, *HSP90AA1* is essential for stabilizing AKT activity, thereby facilitating the phosphorylation and inactivation of FOXO3(*53*) (Fig. 3C). Among divergent genes, 15 and 9 genes were downregulated in the O-A and O-L, respectively (Fig. 3B). Five of these dysregulated genes (i.e., *WWC1, PRKCZ, MAPK4, KSR1, ERBB4)* are involved in enhancing the MAPK cascade activation—a signaling pathway activated downstream of RTKs, similar to the PI3K-AKT pathway (Fig. 3D). *ERBB4* encodes an EGFR, which belongs to RTKs, and was reported previously to be expressed by CTB as a pro-survival factor under hypoxic conditions (*54*). Thus, our findings suggest that in O-A, upregulation of the MAPK cascade might compensate for the downregulation of the PI3K-Akt pathway, promoting downstream target expression, such as Myc targets (Fig. 3A, E). However, this regulatory balance was likely absent in O-L, which could explain the observed downregulation of Myc target genes in O-L (Fig. 3A). Altogether, the PI3K-Akt and MAPK pathways, key downstream effectors of RTK signaling, are dysregulated in obesity and related to fetal overgrowth.

### Transcriptional changes in non-trophoblast cell types due to maternal and fetal growth effects

In VEC, most identified hallmark gene sets were downregulated in both O-A and O-L, including gene sets related to complement signaling, G2M checkpoint, heme metabolism, IL2-STAT5 signaling, KRAS signaling and protein secretion (Fig. 4A). These pathways are related to immune dysregulation, angiogenesis and metabolic stress, which have been shown as placental features of maternal obesity (*55*). Further examining the dysregulated genes within these pathways identified that 49 genes were altered by maternal obesity, and 3 genes were differently regulated in O-A and O-L (Fig. 4B). 9 genes were reported to be critical for regulating VEGFA-VEGFR1 (i.e. FLT1) signaling in angiogenesis (Fig. 4C). Among these, *GNA14, RAPGEF5, NRP1, COL4A2,* and *COBLL1* have been reported to be co-expressed with *VEGFR1* (*55*), while *FN1* and *BMP6* have synergistic effects on VEGFA complex (*56*) (Fig. 4C). The binding of VEGFA complex to VEGFR1 promotes angiogenesis (*57*) and inflammatory cell recruitment (*58*), aligning with the consequence of their dysregulation in maternal obesity. *NRP1* is an adaptor protein forming dimers with VEGFA to bind VEGFR1 (*59*). *FN1* dysregulation has been shown to affect VEGFA expression and relate to cellular apoptosis and autophagy in human umbilical vein endothelial cells (*60, 61*).

**Fig. 4:**
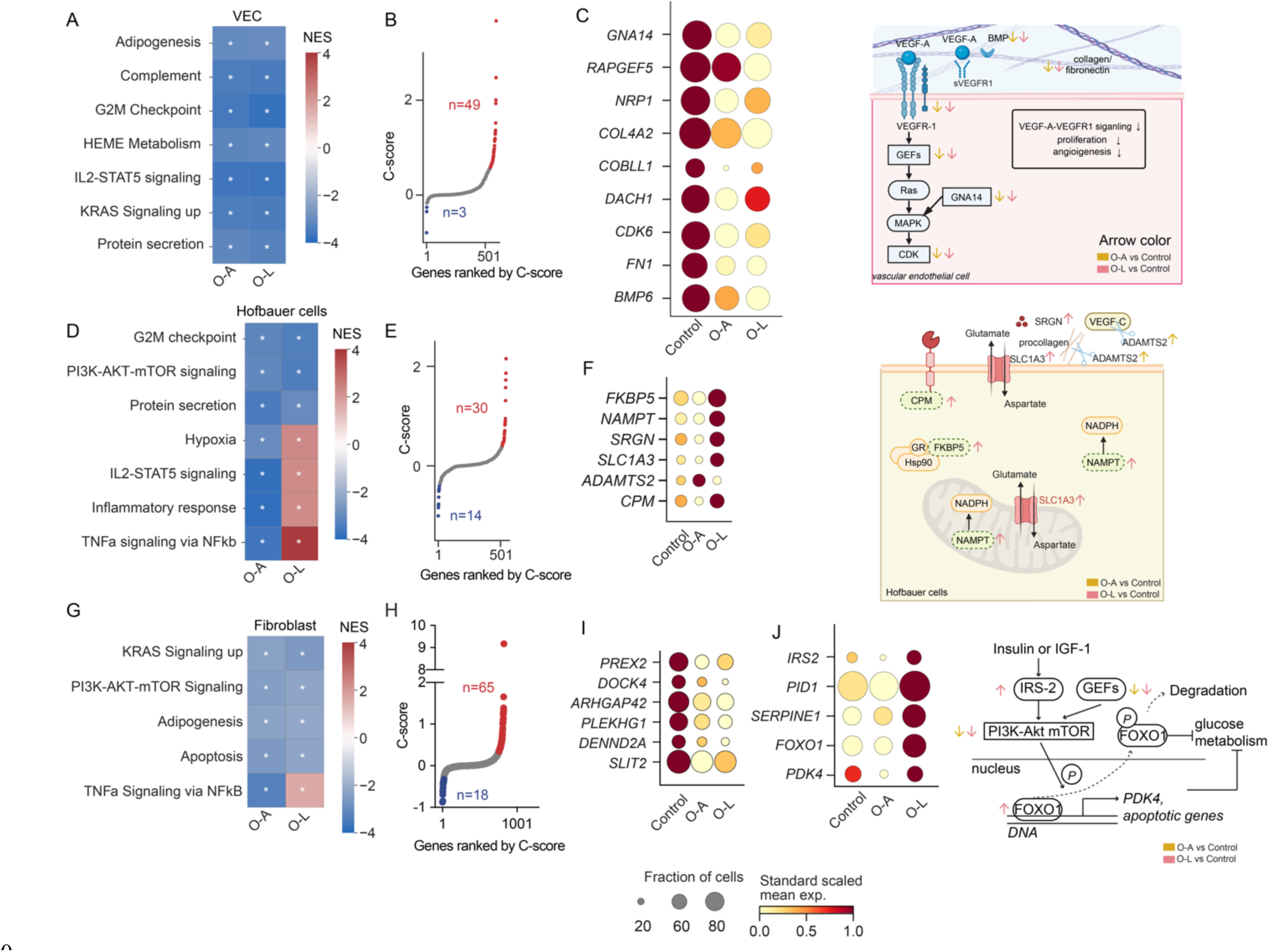
Gene expression changes in vascular endothelial cells, Hofbauer cells and fibroblasts from placentas of mothers with obesity with or without fetal overgrowth. **(A)** Gene set enrichment analysis (GSEA) in vascular endothelial cells (VEC), with the enrichment score of hallmark gene sets for O-A *vs* Control and O-L *vs* Control. **(B)** Scatterplots of genes ordered by C-scores in VEC. **(C)** Dotplot showing expression of genes relevant to FLT1 (VEGFR1) (left). Summary of genes involved in VEGFR1 signaling (right). **(D)** GSEA in Hofbauer cells, with the enrichment score of hallmark gene sets for O-A *vs* Control and O-L *vs* Control. **(E)** Scatterplots of genes ordered by C-scores in Hofbauer cells. **(F)** Dotplot showing expression of genes involved in inflammatory response in Hofbauer cells (left). Summary of these genes in cellular context (right). **(G)** GSEA in fibroblast cells, with the enrichment score of hallmark gene sets for O-A *vs* Control and O-L *vs* Control. **(H)** Scatterplots of genes ordered by C-scores in fibroblasts. (**I**) Dotplot showing expression of genes related to GTPase activity that were changed in maternal obesity, and **(J)** the genes involved in PI3K-Akt pathway and guanine nucleotide exchange factors that were changed according to fetal growth effects. Summary of these genes within the PI3K-Akt-mTOR pathway (right). For **(A)**, **(D)**, **(G)**, NES: normalized enrichment score. *adjusted p-value < 0.05. For (**C**), (**F**), (**I**), the dot size is proportional to the fraction of cells that express the gene, and the color indicating the expression levels standard-scaled between 0 and 1.

In Hofbauer cells, the G2/M checkpoint signaling pathway, PI3K-AKT-mTOR signaling pathway, and protein secretion were suppressed in both obesity groups, suggesting dysregulation of cell proliferation, nutrient-sensing and tissue growth in maternal obesity (Fig. 4D). These changes might represent an adaptive mechanism in Hofbauer cells to curb excessive cytokines and nutrient stress in the placenta under obesity conditions. Notably, hypoxia, IL2-STAT5 signaling, inflammatory response, and TNF-α signaling via NF-kβ were upregulated only in O-L, suggesting that inflammatory and metabolic stress impaired placental homeostasis in O-L (Fig. 4D). Next, we identified 30 genes regulated by maternal obesity (Fig. 4E). Most intriguingly, 14 genes were altered differently in O-L or O-A (Fig. 4F). Among the genes upregulated only in O-L, *FKBP5* encodes for a member of the immunophilin protein family that stabilizes glucocorticoids receptor and promotes NFKβ-mediated inflammation in response to reactive oxidative species (*62*). *NAMPT*, also known as Visfatin or PBEF, has a dual role as an enzyme and a cytokine regulating glucose homeostasis (*63*). *SRGN* encodes a major secreted proteoglycan in macrophages and promotes TNF-α secretion upon lipopolysaccharide stimulation (*64*). Of note, *ADAMTS2* was only upregulated in O-A but not in O-L. *ADAMTS2* is a secreted metalloproteinase that processes procollagen into collagen, a critical step for extracellular matrix stability. Its expression has been implied as part of an anti-inflammatory response to contributing to tissue homeostasis (*65*), which is consistent with increased inflammatory responses in O-L.

Another essential cell type in the villous core is the fibroblast. It is critical for villi architecture, vascular development, immune regulation, and support of trophoblast functions. In fibroblasts, the hallmark gene sets for KRAS signaling, PI3K-AKT mTOR signaling, adipogenesis and apoptosis, were suppressed in both O-A and O-L, while TNF-α signaling via NFKβ was upregulated only in O-L (Fig. 4G). We then identified 65 genes altered by maternal obesity and 18 genes by fetal growth effects (Fig. 4H). Six genes that encode guanine nucleotide exchange factors to regulate GTPase activity were decreased in both O-A and O-L (Fig 4I). These GTPase regulators control multiple small GTPases that are involved in the feedback loop in PI3K activation (*66*). On the contrary, *IRS2*, *PID1*, *SERPINE1, FOXO1, PDK4* were significantly upregulated in O-L while downregulated in O-A (Fig. 4J). Notably, these genes are related to fatty acid response, insulin receptor signaling and RTK signaling. FOXO1 could induce expression of *PDK4* (*67*), and it is also a master regulator for TNF-α induced apoptosis and can inhibit the glucose metabolism (*68*).

### Cell-cell communication regulated by maternal and fetal growth effects in the placenta

Our analyses have so far dissected the gene expression altered by maternal obesity and fetal growth effects in major placental cell types. To further explore whether and how these cell types communicate and coordinate transcriptional regulations in the placenta, we utilized the consensus resources from LIANA+ (*69*) to examine the ligand-receptor (L-R) interactions among cell types. To identify ligands expressed by each cell type that mediate signaling to other cell types, we first constructed a weighted directional regulation network, mapping cell type-L to cell type-R (fig. S3). To evaluate the biological relevance of these interactions, L-R interaction strength was calculated with C-scores of gene pairs weighted by p-values. The altered genes in both O-A and O-L groups were composed the L-R network regulated by maternal effects (Figure. 5A). While altered genes differently between O-A and O-L composed the network related to fetal growth effects (Fig. 5B). This analysis revealed VEC as the main cell type that sent the signals regulated by maternal effects (Fig. 5A). And Hofbauer cells mainly sent signals associated with fetal growth effects (Fig. 5B). Indeed, ligands from VEC, including *FN1*, *COL4A1* and *BMP6,* were among those highly ranked out-degree genes in the network regulated by maternal effects (Fig. 5C). While ligands from Hofbauer cells, including *SPP1*, *PDGFC* and *MAML2*, were highly ranked out-degree genes in the network related to fetal growth effects. Similarly, we then analyzed in-degree scores across cell types, representing the receiving of signaling based on receptor expression (Fig. 5D, E). We identified STB-c as the primary signaling receiver, followed by CTBs, in network regulated by maternal obesity (Fig. 5D). And fibroblasts were the main cell type receiving signaling related to fetal growth from other cell types (Fig. 5E). By ranking in-degree scores of genes across cell types, we identified that integrin complexes (*ITGA3-ITGB1*, *ITGAV-ITGB8*, *ITGA6-ITGB4*) were highly ranked receptors expressed in STB-c regulated by maternal obesity, while *DSCAM* was the highest-ranked receptor expressed in fibroblasts related to fetal growth (Fig. 5F).

**Fig. 5:**
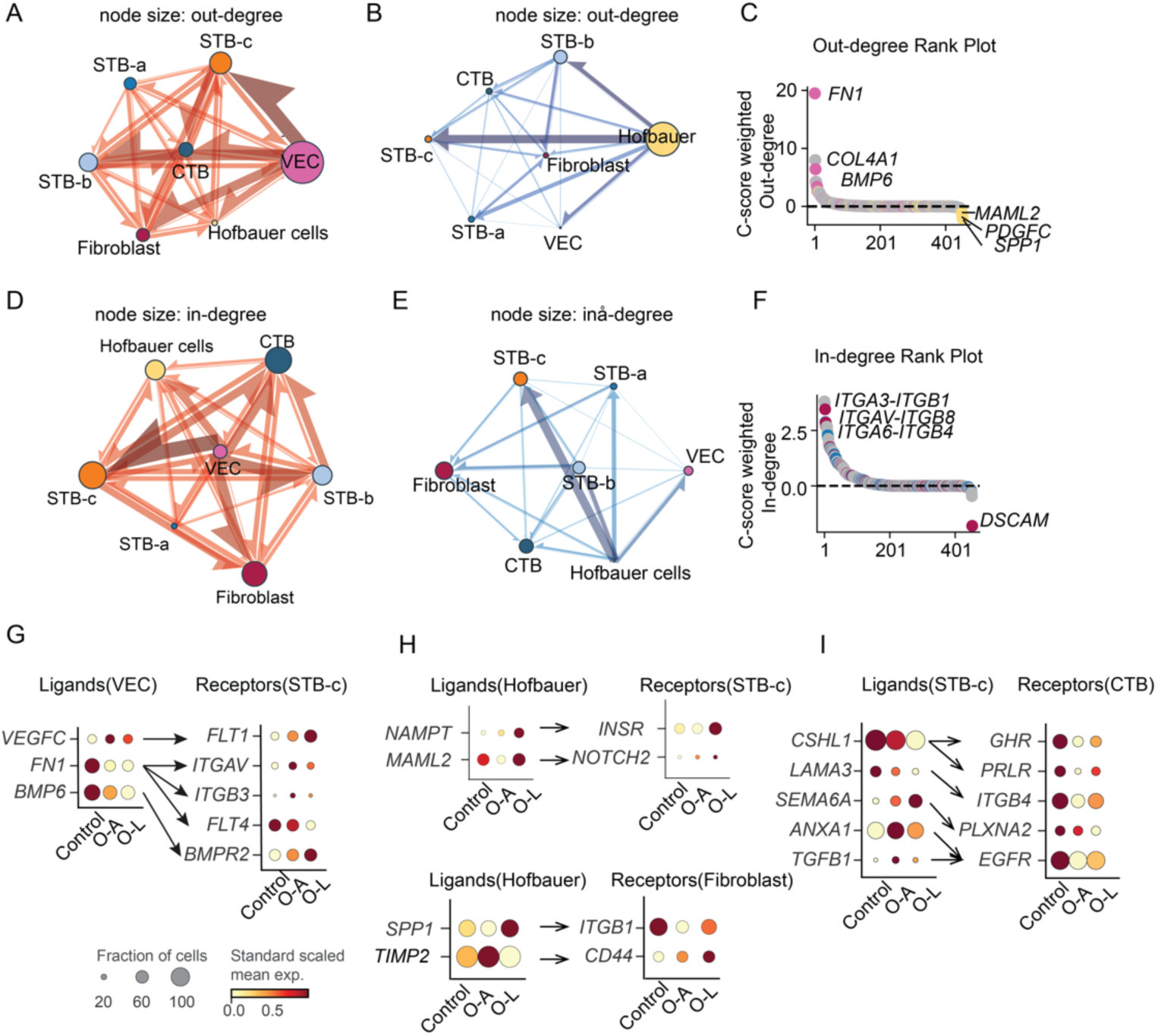
Dysregulated cell-cell communication networks of mothers with obesity, with or without fetal overgrowth. **(A)** Connectome ring plots for cell-cell communication built from differentially expressed ligand-receptor pairs shared in maternal obesity groups and (**B**) differently according to fetal growth effects. Different colors represent different cell types, node size indicating out-degree, and edge direction goes from the sender cell type to the receiver cell type. **(C)** Out-degree analysis indicating the major or sender cells of ligands regulated commonly in maternal obesity or differently according to fetal growth effects. Pink dots represent the vascular endothelial cells, and the yellow dots represent the Hofbauer cells. **(D)** Connectome ring plots built from ligands regulated in maternal obesity and (**E**) differently according to fetal growth effects among major cell types in placental villi from mothers with obesity. Different colors represent different cell types, node size indicating in-degree, and edge direction goes from the sender cell type to the receiver cell type. **(F)** In-degree analysis indicating the major receiver cells with receptors. Red dots represent fibroblasts, and blue dots represent STB-c. **(G)** Dot plot showing the expression of ligands from the VEC)and receptors from STB-c. **(H)** Dot plot showing the expression of ligands from Hofbauer cells and receptors from fibroblasts and STB-c. **(I)** Dot plot showing the expression of the ligands from STB-c and the receptors from CTB.

Specifically, the main cell-cell communication regulated by maternal obesity was observed between VEC and STB-c. The altered L-R pairs such as *VEGFC-FLT1*, *FN1-ITGAV*, -*ITGB3, and -FLT4*, and *BMP6-BMPR2* (Fig. 5G) play a role in vascular formation, tissue structure, and cell fate decisions. Moreover, top cell-cell communications related to fetal growth were observed between Hofbauer and STB-c. The altered L-R pairs included *NAMPT-INSR* and *MAML2-NOTCH2. NAMPT* functions as an intra-NAD biosynthetic enzyme and extracellular inflammation mediator, and elevated extracellular NAMPT was reported in obesity (*70*). Additionally, *SPP1-ITGB1* and *TIMP2-CD44* between Hofbauer and fibroblasts were altered differently in O-A or O-L (Fig. 5H). *SPP1* encodes a multifunctional glycoprotein and the *Spp1*^+^ macrophages emerged under obese conditions in mice (*71*). *TIMP2* signaling was shown to stimulate fibroblast proliferation (*71*).

CTB was also a prominent signaling receiver at vasculo-syncytial membranes. Interestingly, we found that most L-R interactions occurred between STB-c and CTB, involving growth-promoting signaling pairs such as *CSHL1-GHR/PRLR*, *LAMA3-ITGB4*, *SEMA6A-PLXNA2* and *ANXA1/TGFB1-EGFR,* among others (Fig. 5I). *CSHL1* encodes a human placental lactogen, that can bind to the growth hormone receptor consisting of GHR and/or PRLR dimers. Additionally, epidermal growth factor (EGF) is required for proper control of growth and differentiation of CTB into STB (*72*) and altered EGFR expression has been associated with placental pathologies, including intrauterine growth restriction (*73*). Interestingly, receptors for these growth-promoting ligands were decreased in CTBs, suggesting active adaptation to maternal obesity.

### Modeling the obese uterine milieu by co-culturing adipose spheroids and trophoblast organoids by a novel microfluidic system

Although it is well-established that maternal obesity affects placental function and offspring outcomes, the role of different maternal organs and tissue–tissue communication with the placenta remains unclear. Therefore, we investigated whether any observed transcriptional changes in the placenta could be attributed to communication between the adipose tissue and the placenta. To do so, we adopted a microfluidic co-culture system with medium flowing from primary adipose spheroids (ASs) to human term placenta-derived trophoblast organoids (TOs). The main component of the fluidic device contains interconnected tissue chambers, arranged in pairs, enabling parallel perfusion for triplicate experiments (*74*). To ensure gas permeability and conformal contact, a PDMS layer—never in direct contact with the culture media—was applied over the fluidic layer, and finally, an additional layer of PMMA was mounted on the top. It provided a stable chip-to-world connection and ensured sealed perfusion when inlet and outlet connectors are assembled (Fig. 6A). The medium was optimized for glucose, insulin and free fatty acids to simulate an obesity-like environment, consistent with its prior use for ASs culture (*75*). The first chamber pair of the microfluidic chip was designated for a medium control condition without ASs and in the second chamber pair, we placed ASs grown in the “obesity-like” conditions (*75, 76*) before seeded into the device for a 4.5-day co-culture. The ASs were established from white adipose tissue of female origin and cultured for a total of 17 days in “obesity-like” conditions before introduced in the microfluidic device. ASs displayed large lipid droplets stained by BODIPY, consistent with previous (*75*) (Fig. 6B). TOs exhibited a typical morphology with CTBs forming the outer layer shown by E-cadherin staining, and STBs embedded inside the inner layers shown by CGβ staining as shown previously in Matrigel embedded cultures (*77*) (Fig. 6B).

**Fig. 6:**
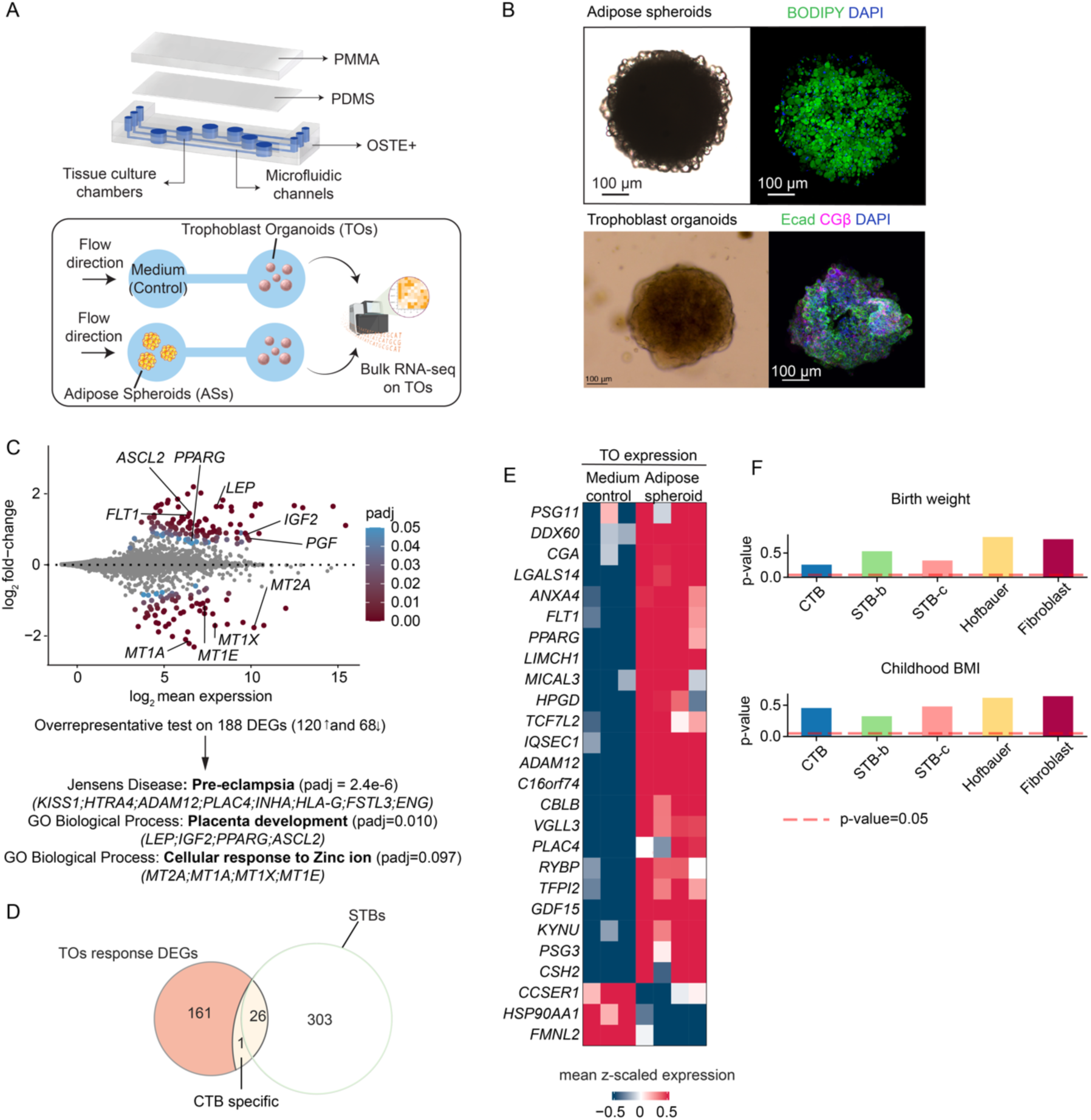
Trophoblast organoids co-cultured with adipose spheroids recapitulated the transcriptional changes observed in STB of mothers with obesity. **(A)** Design of the microfluid device and co-culture experiment of Trophoblast Organoids (TOs) and adipose spheroids (ASs). **(B)** Representative images of ASs (up; bright-field image and confocal image stained with BODIPY and DAPI to highlight neutral lipids and nuclei, respectively) and TOs (bottom; bright-field image and confocal image stained with E-cadherin, CGβ and DAPI to highlight CTBs, STB and nuclei, respectively). Images were captured with 10x magnification. **(C)** MA plot of gene expression in TOs co-cultured with ASs compared with TOs cultured without ASs. **(D)** Venn diagram showing the overlap of the differentially expressed genes in the microfluidic co-culture experiment and trophoblast in maternal obesity groups. **(E)** Expression heatmap for the 26 overlapping genes in (D). (**F**) Enrichment between cell type–specific differentially expressed genes related to fetal growth effects and genomic variants implicated by GWAS for birth weight and childhood BMI computed with MAGMA (*80, 81*).

To examine how TOs were affected by secreted factor from “obesity-like” ASs, we performed bulk RNAseq of the TOs. Indeed, after the continuous exposure to obese ASs, the TO transcriptome was altered, with 188 differentially expressed genes (DEGs; 120 upregulated and 68 downregulated) when compared to medium control condition. Among the upregulated genes, *LEP*, *IGF2*, *PPARG* and *ASCL2* are related to placenta development, while *KISS1*, *HTRA4*, *ADAM12*, *PLAC4*, *INHA*, *HLA-G*, *FSTL3*, *ENG* are related to pre-eclampsia (Fig. 6C). On the contrary, metallothionein-encoding genes *MT1A, MT1E, MT1X,* and *MT2A* were significantly downregulated, suggesting impaired cellular response to ions such as zinc (Fig. 6C). Next, we compared 188 DEGs found in TOs with DEGs of all trophoblast cell types from our snRNA-seq analysis (Appendix Table S4). A total of 27 overlapping genes were identified, with 26 shared with STBs and one shared exclusively with CTB (Fig. 6D), which aligned with the cellular composition of TOs consisting predominantly of STB and a smaller population of CTBs. Importantly, many genes were highly expressed in TOs co-cultured with AS in “obesity-like” conditions (Fig. 6E). These included genes encoding placenta-specific secreted proteins (*46*), including *CGA*, *PSG3, PSG11*, *CSH2* and *TFPI2* (Fig. 6E). *LGALS14*, important for membrane integrity, was also upregulated (Fig. 6E). Interestingly, these 26 overlapping DEGs were all regulated in STB in O-A and O-L from the snRNA-seq analysis. The altered gene expression related with fetal growth effects, however, was not recapitulated by adipose tissue-derived signaling, and could potentially be ascribed to placental-fetal interactions.

To further confirm that, we investigated whether the identified fetal growth effects are linked to genetic loci, previously identified by genome-wide association studies (GWAS), associated with birth weight (*78*) and childhood BMI (*79*). We used MAGMA, a tool for generalized gene-set analysis of GWAS data, to analyze the statistical association between the aggregate genetic risk at the gene-level (*80, 81*). To ensure reliable results, we excluded cell types from this analysis with less than 10 DEGs. For remaining placental cell types, we did not find strong associations between genes that were altered related to fetal growth effects at the transcription level and risk genes at the genetic variation level (Fig. 6F). This suggests that transcriptional alterations identified as fetal-growth effects might not be attributed to fetal genetic effects but placental-fetal interactions to accommodate the fetal growth.

## Discussion

Maternal obesity during pregnancy is associated with several adverse outcomes for both mothers and offspring. A deeper deciphering of the molecular mechanisms underlying placental responses is crucial for future interventions and therapeutic development. In this study, we examined placentas of women with obesity who did not develop other pregnancy complications but delivered either AGA or LGA infants (*28*). This could help minimize confounding factors and allow for a clearer understanding of inherent placental alterations. Given that the human placenta is a highly specialized organ with distinct cell types, including the multinucleated STB layer, we utilized snRNA-seq to reveal cell-type-specific transcriptional responses to maternal obesity and fetal growth. Notably, we were able to identify a set of transcriptional changes across placentas affected by maternal obesity, regardless of fetal growth effects, highlighting these as maternal effects. Moreover, we identified a distinct set of transcriptional changes that differed between O-A and O-L, potentially related to different fetal growth effects. We further examined cell-cell communication in mediating these maternal or fetal growth effects, highlighting key L-R pairs. Notably, VEC, STB, Hofbauer cells and fibroblasts played prominent roles in such communications. These findings are summarized in Figure 7.

**Fig. 7:**
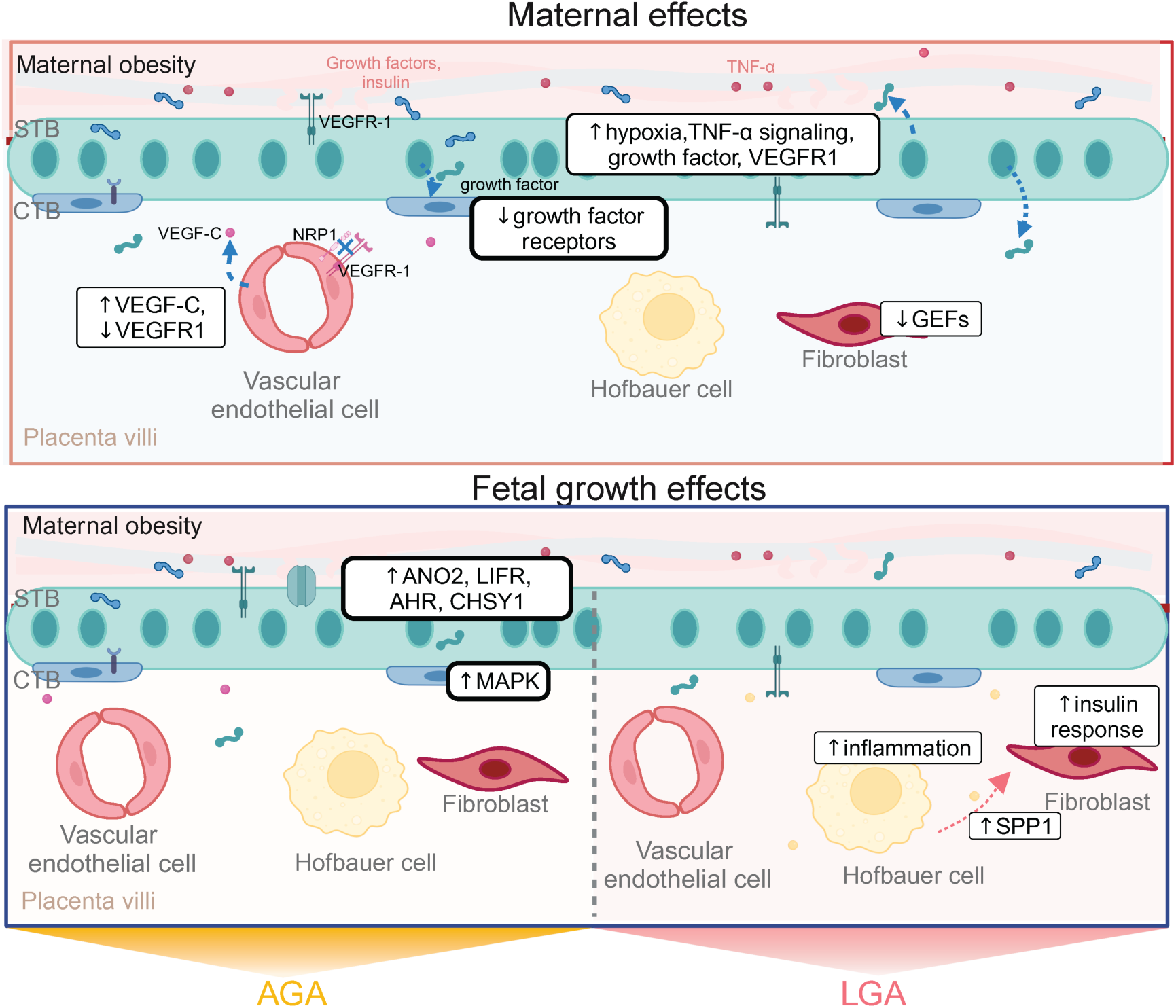
Schematic summary showing that gene expression changes in maternal obesity and different gene expression changes according to fetal growth effects, highlighting the genes involved in each cell types.

Previous studies have shown that altered metabolites, hormones and cytokines - such as IL-6 and TNF-α - in maternal obesity can be sensed by trophoblast cells, thereby influencing trophoblast proliferation, nutrient uptake and transport (*82, 83*). Furthermore, activation of phospholipase A2 by TNF-α and leptin in human trophoblast cells has been proposed as a mechanism contributing to excessive fetal fat accumulation and neonatal adiposity (*84*). In line with this, our findings reveal that STB exhibit an enrichment of pathways related to elevated hypoxia and TNF-α signaling via NF-kβ in maternal obesity. These responses were potentially driven by alterations in genes involved in the cross-regulations of these pathways, such as *FOS*, *FOXO1*, and *SLC2A1*. *SLC2A1* is of particular interest as it encodes GLUT1 to promote glucose uptake, suggesting that, similar to other metabolic organs, there is an interplay between hypoxia, inflammation and excessive nutrient transport specifically in the STB. Interestingly, we have recently demonstrated how mild hyperglycemia can still induce placental hypoxia in mice, suggesting excessive hypoxia as a common placental signature in maternal metabolic disorders (*85*).

Notably, in maternal obesity, CTBs downregulated genes involved in PI3K-Akt signaling pathways, particularly those RTKs associated with the *GHR*, *PRLR*, and *CSF3R*. It is likely that the altered expression of these receptors in CTBs is an active adaptation to the altered ligands expressed in STB. When analyzing these L-R interactions between two cell types within placental villi, we further found that mature STB-c altered the expression of growth-promoting ligands that could bind to receptors in CTBs, which were generally downregulated, including growth factors receptors binding to insulin-like growth factor, prolactin, and colony-stimulating factors (*86, 87*). Likely, downregulation of these receptors in CTB followed by PI3K-Akt signaling pathways could dampen the tissue growth signaling and serve as a potential link to maternal obesity.

Importantly, we further uncovered the molecular regulation related with the fetal growth in maternal obesity. Notably, TNF-α signaling and inflammatory response were decreased in the Hofbauer cells in O-A, whereas they were prominently increased in O-L, implying that LGA in maternal obesity is strongly associated with inflammation. Aligning with this, *NAMPT* and *SLC1A3,* which play a dual role in this inflammatory response and metabolism (*88*), were exclusively upregulated in O-L. These genes likely facilitate the immune-metabolic modulation by Hofbauer cells related to fetal overgrowth. Moreover, L-R network analysis revealed that Hofbauer cells sent the ligands differently altered by fetal growth effects to other cell types in the villi. Hofbauer cells in O-L highly expressed the ligand *SPP1,* and SPP1^+^ macrophages have been reported to be in proximity to hypoxic areas in tumors, to support tumor progression by promoting cell survival, angiogenesis, and immunosuppression (*89*). In addition, dysregulated Hofbauer polarization and immune regulation can also restrict fetal growth (*90–92*). Therefore, Hofbauer cells may serve as a crucial link in modulating fetal growth effects in response to hypoxic and inflammatory uterine environment in maternal obesity.

Moreover, an increase in TNF-α signaling in fibroblasts was also observed exclusively in O-L. This signaling may impair insulin action on fibroblasts by inhibiting the activation of IGF-1/insulin hybrid receptors (*93*). In fibroblasts, the concurrent increase of TNF-α signaling and genes involved in insulin receptor signaling, such as *IRS2,* suggests that these regulatory mechanisms may be another contributor to fetal overgrowth. In line with this, we identified fibroblasts as the major signaling receiver associated with fetal overgrowth, with the most expressed ligand being macrophage-derived *SPP1*, which has been shown to activate myofibroblasts in fibrosis (*94*). Fibroblasts and stromal cells in the term placenta have been understudied in prior research, our analysis highlights their transcriptional responses to maternal obesity, particularly with fetal overgrowth. Interestingly, TNF-α signaling was increased in STB of both obesity groups but only increased in Hofbauer cells and fibroblasts in O-L. This may indicate that the STB response to TNF-α is predominantly shaped by direct exposure to maternal circulating inflammatory signals, whilst within the placental villi, these maternal signals need to be dampened to achieve appropriate fetal growth. Regulatory mechanisms to protect the placental environment from excessive inflammatory responses might be critical to proper fetal growth, while excessive inflammation within villous niche likely related to fetal overgrowth. It would be valuable to further explore whether targeting Hofbauer-specific inflammatory signaling pathways can mitigate fetal overgrowth and thereby reduce the linked risk of cardiometabolic disorders in adulthood.

When investigating the intercellular communication network, we found that STB is the primary ‘receiver’ for the overall L-R pairs in maternal obesity regardless of fetal growth. Whether these signals originate from maternal circulation or villous cells cannot be concluded from the snRNA-seq data. Complement to this, we used a novel microfluidic system to co-culture TOs and ASs. This confirmed that certain signals were likely of maternal origin, due to tissue-tissue communication. Our microfluidic co-culture system displayed several advantages for studying the functional impact of tissue communication. The unidirectional media flow from ASs to TOs simulated the effects of potential metabolites and cytokines from the maternal circulation on the placenta. As DEGs in TOs exhibited a greater overlap of altered genesto maternal obesity in STB, including *FLT1* and *PPARG,* compared to other cell types, this further suggests the key role of STB regulation by the maternal environment. Our co-culture system could not find any genes altered differently by fetal growth effects in STBs, suggesting that factors other than the uterine milieu governs such fetal effects. Interestingly, genes transcriptionally altered differently by fetal growth effects are not among those genetic loci identified by GWAS of birthweight and childhood obesity. This further suggested the importance of the placental-fetal interplay. Future investigations into the intricate dynamics of placental–fetal interactions will fundamentally reshape our understanding of this developmental processes. However, such experiments require innovative technological advances on embryo models. Importantly, our findings reveal promising molecular targets for further exploration.

In conclusion, our study demonstrates an intricated cell type-specific regulatory network within the placenta, driven by critical pathways such as hypoxia, nutrient transport, inflammation, and TNF-α signaling in maternal obesity and fetal overgrowth. Given the rising prevalence of global obesity, these mechanistic insights will deepen our understanding of placental interplay with fetal development. By precisely modulating these pathways, we can potentially mitigate pregnancy complications and lower the long-term risk of adverse health outcomes in offspring.

## Materials and Methods

### Experimental Design

The Institutional Ethics Committee for Research on Humans of the Faculty of Medicine, University of Chile (Protocol Nr. 033-2013, 201-201, 236–2020) reviewed and approved the protocol. At the time of recruitment, each participant received all information regarding the study, voluntarily enrolled and signed the informed consent. Women were recruited before delivery at the Maternity Service of the Hospital San Juan Dios, West Health Division, Santiago, Chile between 2014 and 2022. From the 328 placentas collected during this period, women with pregnancy complications such as hypertension, preeclampsia, thyroid disorders, GDM, preterm delivery, a fetus with malformations or chromosomal aberrations, and those with pregestational antecedents of diabetes, polycystic ovary syndrome, infertility, and assisted fertilization (IVF or ICSI) as well as those who declared to smoke, drink alcohol or take drugs during the pregnancy were excluded from this study. To the purpose of this study, women with normal pregestational weight (BMI between 18.50 -24.99 kg/m^2^, n = 22) or pregestational obesity grade I (BMI between 30 – 35.7 kg/ m^2^, n = 25) aged between 18 and 34 years old, that delivered from vaginal deliveries were further selected. Birth weight appropriate to gestational age (AGA) was defined as a birthweight more than 10th and below 90th percentile and birth weight large to gestational age (LGA) was defined as a birthweight above 90th percentile of the sex-specific and gestational age-specific reference mean(*6*). For single-nucleus analysis, placenta samples from control (normal-weight mothers and AGA babies, n=4), O-A (mothers with obesity and AGA babies, n=4), and O-L (mothers with obesity and LGA babies, n=4) were selected. And the maternal age, height, gestational age, and baby sex were not different between groups.

### Sample collection

After delivery, placentas were collected as previously described (*28*) and snap frozen immediately in liquid nitrogen. Briefly, placentas from full-term pregnancies were collected immediately after delivery and processed within 30min. Each placenta was sectioned transversely using a sterile scalpel near the cord insertion site (approximately 5 cm) and villous part was used for snRNA-seq. The samples were then conserved in -80°C freezers until transport to Sweden when they were kept on dry ice and then store again in -80°C freezers. Newborn data such as sex, weight, height, and head circumference were recorded. Placental efficiency was calculated as the ratio of neonatal weight (g) to placental weight (g).

### Nucleus isolation

Protocol for nucleus isolation was adapted from the protocol optimized for human placenta samples by Ludivine Doridot and her team (Institut Cochin, France). Briefly, 30mg of frozen placenta was chopped into small pieces on dry ice and added to douncer with cold NP-40 lysis buffer containing RNase inhibitor. After 10min of incubation on ice, lysis buffer was diluted in wash medium (cold PBS with 2% BSA and RNase inhibitor) and 10 strokes with the loose pestle were performed. Samples were further filtered (100µm) and centrifuged at 600g, for 5 min at 4C. Pellet was resuspended in cold wash medium, and then further filter (40µm) and centrifuged again. After resuspension in nuclei resuspension buffer (1% BSA and RNase inhibitor solution in DPBS), the pellet was filtered one last time (20µm).

The number of isolated nuclei was determined using trypan blue staining and automated cell counter (Biorad, TC20) and quality of nuclei was assessed by visualization under microscope (EVOS XL Core, ThermoFisher Scientific).

### 10x Genomics library preparation and sequencing

Suspension of good quality nuclei were processed as manufacturer’s protocol for the Chromium Single Cell 3’ kit v3.1 (10X Genomics). Library preparation was performed to obtain between 1,000 and 10,000 nuclei per reaction. Libraries were then sequenced to obtained 90GB of data (paired end, 150bp) per reaction on the Illumina NovaSeq 6000 at Novogene, UK.

### snRNA-seq processing

Gene counts were obtained by aligning reads to the GRCh38 genome using Cell Ranger software (10x Genomics). We used scanpy (*95*) to process and cluster the expression profiles and infer cell identities of major cell classes. Cellbender (*96*) was used to remove background noise. In addition, using Scrublet (*97*), cells that were labeled as doublets were removed.

For the UMAP visualization of individual major cell type classes, the analytical Pearson residue was used to find the highly variable genes from individual batches. We selected the set of relevant principal components based on elbow plots. High resolution cell clusters were identified using the Leiden clustering algorithm. We annotated cell types using previously published marker genes and single-cell RNA-sequencing data (*29*).

For the Pseudo-time analysis of trophoblast nuclei, Slingshot (*98*) was applied. Slingshot was designed to identify a minimum spanning tree (MST). In this setting, pCTB was assigned as an initial cell state and STB was classified as a terminal cell state.

We integrate the syncytiotrophoblast data generated in this study with the syncytiotrophoblast data from the public human healthy term placenta snRNA-seq data (*99*) into the same embedding using scANVI (*38*). We transferred their label onto our nuclei and compared the transferred labels with our labels using the sihouette metric. The silhouette metric’ values range from [−1, 1] with 1 indicates distinct labels, 0 indicates overlapping labels. Thus, the absolute value of the silhouette width is used to measure how well labels are mixed.

We applied a Bayesian model in scCODA (*100*) for testing the compositional differences across groups. pCTB population was selected as the reference cell type, which is assumed to be unchanged in absolute abundance, and parameters inference was calculated via Hamiltonian Monte Carlo (HMC) sampling.

### Differential expression of obesity groups *vs* control

We performed differential expression analyses using a Wald test with a negative binomial regression model using the formula “∼ 1 + group + sex + log_number_of_genes” on raw counts. Such analysis is deployed in R Seurat package (*101*). By default, in FindMarkers function in Seurat, genes with more than one count in at least 1% of cells were considered. This rendered 6,000 to 7,000 genes to be analyzed. FDR was estimated with Benjamini-Hochberg method. The statistics from this analysis were used as the input in the following method section.

For the gene set enrichment analysis, we ranked the genes in O-A vs. Control and O-L vs. Control differential gene expression analysis respectively by fold changes. We then performed GSEA (*102*) to get the normalized enrichment scores. Specifically, an enrichment score was calculated by walking down the list of genes, increasing a running-sum statistic when a feature in each hallmark gene set is encountered and decreasing it when it is not. The final score is the maximum deviation from zero encountered in the random walk. Finally, a normalized score was obtained by computing the z-score of the estimate compared to a null distribution obtained from N random permutations. The p-values were adjusted by the BH method (*103*) to control the false discovery rate.

Gene Ontology enrichment analyses were performed using Enrichr (*55*). Specifically, Fisher’s exact test was used to test the overrepresentation of the gene sets of interest from this study in known gene sets from MsigDB Hallmark database. The p-value from the test was adjusted with the BH method.

### Scoring the shared or different DEGs between comparisons

To measure the similarities or differences in differential gene expression between the two obese groups and the normal weight group, we construct a score based on the fold changes of the genes and the false discovery rates (FDRs) of the two comparisons:

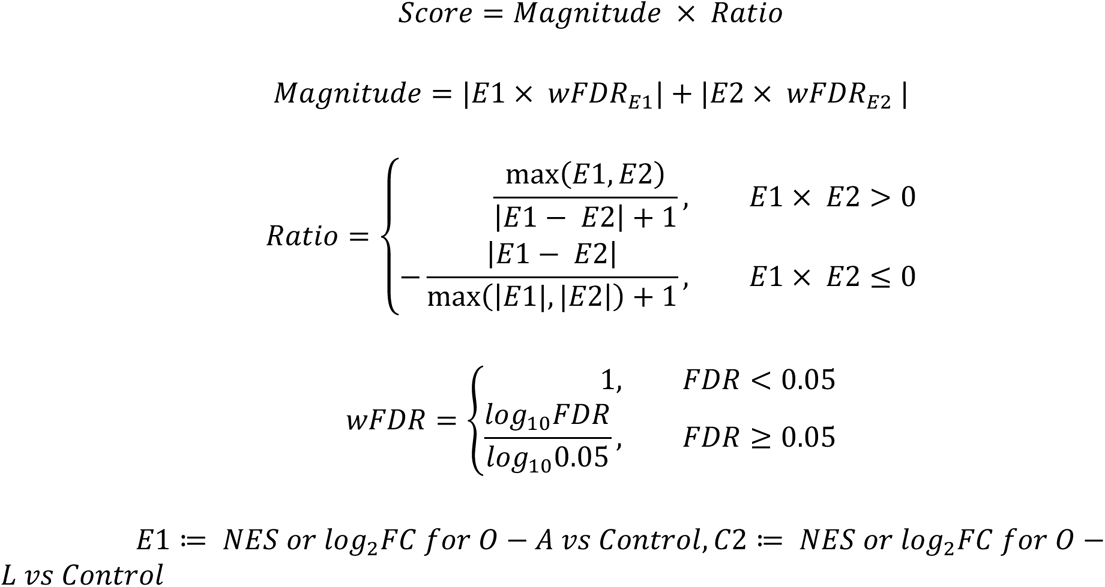

A positive score indicates that the genes were both increased and decreased in the two comparisons. The higher the score, the more similar and larger fold changes were. Conversely, a negative score indicates that the gene was oppositely expressed in the two obese groups compared to the normal weight group. This means that it was increased in one comparison group and decreased in the other. The more negative the score, the more divergent and larger fold changes the gene had. In this algorithm, genes with zero-fold changes were scored as zero.

To measure the significance of the scores of genes that were differentially expressed (score≠0), we performed a permutation test. The fold changes and FDRs of all these genes in each comparison were permuted 40,000 times if the number of genes with a score≠0 was over 200, otherwise, they were permuted (number of the genes)^#^ times. The p-values were calculated to the observed scores as the frequencies of the permutated scores > the observed score when the observed score is positive or as frequencies of permuted scores < the observed score when the observed score is negative. A one-sided p-value < 0.05 was used for significance.

### Cell-cell communication (CCC) network analysis

Ligand–receptor interactions contrasts across cell type pairs were inferred using the consensus ligand–receptor database from LIANA+ (*104*). The ligand-receptor edges were weighted by the sum of C-scores for the corresponding genes obtained with the method stated in the preceding section. This generated the cellA–cellB–ligand–receptor interaction tetramers weighted by the similarity (C-scores) of the response in the two obesity groups vs control. The mean p-values for the C-scores of the corresponding genes were assigned to the tetramers indicating significance of the interaction as a response to maternal obesity.

From the weighted CCC network, the in-degree, out-degree, and betweenness centrality were calculated. Betweenness centrality measures the extent to which a vertex lies on paths between other vertices. Vertices with high betweenness may have considerable influence within a network by virtue of their control over information passing between others. They are also the ones whose removal from the network will most disrupt communications between other vertices because they lie on the largest number of paths taken by messages.

### Adipose spheroid establishment

Adipose spheroid (AS) obesity conditions were established as previously described. Briefly, cryopreserved human primary stromal vascular fraction of white adipose tissue of female origin were seeded in culture flasks with DMEM/F-12 Glutamax, 10% FBS and 1% penicillin-streptomycin. After reaching the confluency of around 70%, cells were trypsinized and seeded into ultra-low attachment 96-well plates (Corning) with 5,000 cells per well and centrifuged at 150g for 2 min. After 4 days, the culture medium was changed to serum-free differentiation medium (William’s E supplemented with 2mM L-Glutamine, 100 units/ml penicillin, 100µg/ml streptomycin, 10µg/ml insulin, 10µg/ml transferrin, 6.7ng/ml sodium selenite and 100nM dexamethasone; *differentiation cocktail*, 500µM 3-isobutyl-1-methylxanthine (IBMX), 10nM hydrocortisone, 2nM 3,3’,5-Triiodo-L-thyronine, 10μM rosiglitazone, 33μM biotin and 17μM pantothenic acid, and *free fatty acids (FFAs)*, 160µM oleic acid; 160µM palmitic acid conjugated to 10%BSA at a molar ratio of 1:5). After 48h, half of the medium was changed and subsequently every 3-4 days for 17 days. After differentiation, the spheroids were transferred into the maintenance media supplemented with FFAs for four days before the chip co-culture and the media was exchanged every 2 days to phase out the differentiation growth factors.

### Trophoblast organoid culture establishment

Villous trophoblast primary cells were isolated from human placental tissue as described previously (*77*). Briefly, the isolated villi fragmented and washed intensely. Then, the tissue was digested with 0.2% trypsin (Alfa Aesar, J63993-09)/0.02% EDTA (Sigma-Aldrich, E9884) and 1 mg/mL collagenase V (Sigma, C9263), and further disrupted by pipetting up and down with a serological 10mL pipette, around 10-15 times. The cellular digests were combined and washed with Advanced DMEM/F12 medium (Life Technologies 12634-010), and cells were counted with a automatic cell counter (TC20 BioRad). After, the cells were pelleted and resuspended in pre-thawed Matrigel (Corning 35623). The volume of Matrigel is dependent on cell counting and on pellet size; in general, at least 10x of cell pellet volume should be used. 20µL domes of Matrigel/cells were seeded in a 48-well culture plate and placed in an incubator at 37°C with 5% CO_2_ for 3 minutes, then the plate was turned upside down so cells distribute evenly within the Matrigel domes. The domes were incubated for a total of 15 minutes before supplemented with 250µL of trophoblast organoid maintenance medium, composed with: Advanced DMEM/ F12 (Life Technologies, 12634-010); 1x B27 (Life Technologies, 17504-044); 1x N2 (Life Technologies, 17502-048); 10% FBS (Cytiva HyClone, SH30070.03); 2mM L-glutamine (Life Technologies, 35050-061); 100μg/mL Primocin (InvivoGen, antpm-1); 1,25mM NAC (SigmaAldrich, A9165), 500nM A83-01 (Tocris, 2939); 1,5μM CHIR99021 (Tocris, 4423); 50ng/mL hEGF (Gibco, PHG0314); 80g/mL human R-spondin1 (R&D systems, 4645-RS-100); 100ng/mL hFGF2 (Peprotech, 100-18C); 50ng/mL hHGF (Peprotech, 100-39); 10mM nicotinamide (Sigma-Aldrich, N0636-100G); 5μM Y-27632 (Sigma, Y0503-1MG); 2.5μM prostaglandin E2 (PGE2, R&D systems, 22-961-0).Medium was replaced every 2-3 days. hCG was measured to ensure proliferation of trophoblast cells using professional pregnancy test strips (Gravidetstest GI29100, Medistore). Small trophoblast organoids became noticeable and positive hCG tests were obtained around 10-15 days after establishment.

### Maintenance and recovery of trophoblast organoids

Passages were performed every 5-7 days, as previously described by (*77*). Each Matrigel dome was disintegrated by pipetting the medium up and down and by scraping. Organoid suspension was washed and digested in pre-warmed StemPro Accutase (Gibco, A11105-01) with Y-27632 (Sigma, Y0503-1MG) for 10 minutes at 37°C. After carefully removing the supernatant, 200µL of Advanced DMEM/ F12 (Life Technologies, 12634-010) was added before performing mechanical dissociation by at least 200 ups and downs, using an automatic pipette. Cell suspension was washed and resuspended in cold Matrigel and plated into 20 µL domes with desired cell density. 250µL of trophoblast organoid medium was added, after 15 minutes of incubation at 37°C with 5% CO_2._

After 7 days of culture, TOs were removed from Matrigel. For that, 250µL of Cell Recovery Solution (Corning 354253) was added to each well. The plate was incubated on ice for 45 minutes. The Matrigel domes were dissociated by gently pipetting up and down, the organoid/cell recovery solution was collected and centrifuged. After removing the supernatant, the organoid pellet was washed and resuspended in cold pre-thawed Matrigel. Tubes were kept on ice until the assembly of the microfluidic co-culture.

### Co-culture of trophoblast organoids with adipose spheroids on a microfluidic device

To co-culture obese AS with TOs, we used multilayered microfluidic devices made of off-stoichiometric thiol–ene–epoxy (OSTE+), poly(methyl methacrylate) (PMMA) and polydimethylsiloxane (PDMS). The OSTE+ parts, which hold the tissue chambers (D = 6 mm) and microfluidic channels (W= 1000 µm, H = 500 µm), were produced from OSTEMER-322 (Mercene labs, Sweden), as previously described (*74*). The PDMS layer was cast and the PMMA layer was cut using a micromilling machine (Minitech Machinery Corp., USA). The AS were grown and differentiated for 17 days and washed in maintenance media before the chip culture. Between 20 to 30 AS were loaded in the first chambers of the co-culture chips and 17 µL domes of TOs (grown for seven days after splitting) were seeded in Matrigel in the second chambers of all devices. The control chips were only seeded with TOs. To mimic an obese environment, obesogenic media (previously described) was perfused at a constant flow rate of 5µL/minute using an NE-1600 SyringeSIX Programmable Multichannel Syring Pump (New Era Pump Systems Inc.). After 4 days, TB-Org were collected for RNA-extraction.

### RNA extraction and sequencing of trophoblast organoids

TOs for RNAseq analysis were recovered from Matrigel using Cell Recovery Solution (Corning, 354253), as described previously. 500µL of cold TRI reagent (Sigma-Aldrich) was added and incubated on ice for 5 minutes. Then 100µL of chloroform was added, vortex mixed and centrifuged. Around 200µL of supernatant was recovered and the same volume of isopropanol was then added. The mixture was further mixed and loaded on Minicolumn (ReliaPrep RNA Miniprep, Promega). The next steps were performed following manufacturer instructions and included DNase treatment.

Prime-seq protocol was used to prepare RNA sequencing libraries as previously described (*105*). Briefly, samples with 10ng/µL of extracted RNA were used for reverse transcription and pre-amplification. The cDNA was quantified using the PicoGreen dsDNA Assay Kit (ThermoFisher Scientific) and qualified using the Bioanalyzer High Sensitivity DNA chip (Agilent), before subsequent library construction using the NEBNext Ultra II FS Kit (New England Biolabs). Then the libraries were sequenced on Illumina NovaSeq 6000 at Novogene, UK at an average depth of 20 million reads per sample.

### Downstream analysis of trophoblast organoid RNA sequencing

Quality of the raw data was checked using FastQC (version 0.11.9), trimmed of poly(A) tails using Cutadapt (version 4.1), before filtering using the zUMIs pipeline (version 2.9.7) (*106*). The filtered data was then mapped to the human genome using STAR (version 2.7.10a) (*107*), and reads were counted using RSubread (version 2.12.0) (*108*). Lowly expressed genes (count sum <50 for all samples) were filtered out before differential analysis. Differential expression analysis on RNA sequencing data was done using the DESeq2 package (version 1.36.0) (*109*). Differentially expressed genes (DEGs) were defined as adjusted p-value<0.05 and fold-change>2 or <-2.

### Statistical Analysis

Variables (maternal age, height and BMI, birthweight, placental weight and efficiency) were analyzed using GraphPad Prism 9.4.1 for Windows (GraphPad Software, San Diego, California USA, www.graphpad.com). For cohort comparisons, t test with Welch’s correction were applied. For snRNA-seq sample data, Kruskal-Wallis test followed by two stage linear step procedure of Benjamini, Krieger and Yekutieli for adjustment were performed. For all comparisons, a variable was considered as significantly different between group for p after correction < 0.05. In the text, data are represented as mean ± SEM.

## Acknowledgments

This work is supported by the Swedish Medical Research Council: project nos. 2018-02557(Q.D.), 2020-00253(Q.D.) and 2022-00550 (E.S.-V.), 2021-02801 (V.M.L.), 2023-03015 (V.M.L.) and 2024-03401 (V.M.L.); the Knut and Alice Wallenberg Foundation: 2019.0211 (Q.D.) and VC-2021-0026 (V.M.L.); Karolinska Instiutet faculty funded position (Q.D.); Diabetes foundation: DIA2022-733 (Q.D.); Barndiabetes fonden (Q.D.); Distinguished Investigator Grant – Endocrinology and Metabolism, Novo Nordisk Foundation: NNF22OC0072904 (E.S.-V.); the Diabetes Foundation: DIA2021-633 (E.S.-V.); the Karolinska Institutet-China scholarship council program (H.J.); the Robert Bosch Foundation (V.M.L.), as well as the Innovative Medicines Initiative 2 Joint Undertaking (JU) under grant agreement No. 875510 (V.M.L.). The mapping of the sequencing data and ambient removal were performed using resources provided by the National Academic Infrastructure for Supercomputing in Sweden (NAISS), partially funded by the Swedish Research Council through grant agreement no. 2022-06725.

## Author contributions

H.J contributed to the design of the work, analysis, and interpretation of data, the formulation of the C-score used in this work, and have drafted the work; E.D contributed to the design of the work, and the acquisition of the data; D.P contributed to the acquisition of the trophoblast organoids data and has revised the work; N.T contributed to the acquisition of the adipose spheroids and microfluidic co-culture system; R.Z.S fabricated and characterized the microfluidic devices; M.O contributed the acquisition of the placental tissues; P.R.J contributed to the acquisition of the trophoblast organoids and has revised the work; A.Z contributed to revise the work; C.L contributed to the library construction the placental tissues; M.A.M contributed to the acquisition of the placental samples; E.S-V contributed to the acquisition of the samples and revise the work; V.M.L contributed to microfluidic co-culture system and revise the work; Q.D conceptualized and designed the work, interpretation of the data, drafted the work. All authors contributed to revising the manuscript.

## Disclosure and competing interests statement

V.M.L. is co-founder, CEO and shareholder of HepaPredict AB, as well as co-founder and shareholder of Shanghai Hepo Biotechnology Ltd. The other authors declare no competing interests.

## Data and materials availability

Raw snRNA sequencing data from the placenta tissues and Raw RNA sequencing data from trophoblast organoids were deposited in EGAS50000000834. The explanation and script for C-score calculation and is available at https://github.com/DengLab-KI/cscore. The notebooks to reproduce the presented results is available at https://github.com/DengLab-KI/obesity_paper. Source data for reproducing the figures are deposited at Zenodo https://doi.org/10.5281/zenodo.14549513. Data on birth weight has been contributed by the EGG Consortium using the UK Biobank Resource and has been downloaded from www.egg-consortium.org. Data on childhood body mass index have been contributed by the EGG Consortium and have been downloaded from www.egg-consortium.org.

## Notes

https://doi.org/10.5281/zenodo.15053792

https://github.com/DengLab-KI/cscore

### Competing Interest Statement

Volker M. Lauschke is co-founder, CEO and shareholder of HepaPredict AB, as well as co-founder and shareholder of Shanghai Hepo Biotechnology Ltd. The other authors declare no competing interests.

### Summary of Updates

To better convey biological knowledge, we defined the common responses as maternal effects and the divergent responses as fetal growth effects under maternal obesity condition.

